# Latent factor in Brain RNA-seq studies reflects cell type and clinical heterogeneity

**DOI:** 10.1101/2022.11.13.516360

**Authors:** Rami Al-Ouran, Chaozhong Liu, Linhua Wang, Ying-Wooi Wan, Xiqi Li, Aleksandar Milosavljevic, Joshua M. Shulman, Zhandong Liu

**Affiliations:** Department of Data Science and Artificial Intelligence, Al Hussein Technical University, Amman, Jordan; Jan and Dan Duncan Neurologic Research Institute, Texas Children’s Hospital, Houston, TX 77030, USA; Department of Pediatrics, Baylor College of Medicine, Houston, TX 77030, USA; Department of Molecular and Human Genetics, Baylor College of Medicine, Houston, TX 77030, USA

## Abstract

With the growing availability of Alzheimer’s disease (AD) transcriptomic data, several studies have nominated new therapeutic targets. However, a major challenge is accounting for latent (hidden) factors which affect the discovery of therapeutic targets. Using unsupervised machine learning, we identified a latent factor in brain tissue, and we validated the factor in AD and normal samples, across multiple studies, and different brain tissues. Moreover, significant metabolic differences were observed due to the latent factor. The latent factor was found to reflect cell-type heterogeneity in the brain and after adjusting for it, we were able to identify new biological pathways. The changes observed at both transcriptomic and metabolomic levels support the importance of identifying any latent factors before pursuing downstream analysis to accurately identify biomarkers.

## Introduction

Alzheimer’s disease (AD) is a neurodegenerative disorder affecting more than 5 million Americans, a number that is expected to rise to almost 14 million by 2050 ^1^. AD is characterized by pathological hallmarks, such as amyloid plaques and intraneuronal neurofibrillary tangles, and there are currently no effective disease interventions ^2^. Due to the complexity and heterogeneity of the disease, we have an incomplete understanding of its etiology which hampers precision medicine’s ability to discover valid therapeutic targets for AD.

One promising approach for discovering new potential AD therapeutic targets is to data mine high-throughput transcriptome profiles of human brains. The Accelerating Medicines Partnership-Alzheimer’s Disease (AMP-AD) Target Discovery Project has profiled ~2,000 human brain autopsy samples (bulk-RNAseq). Initial analysis by the AMP-AD consortium has discovered several new biological networks and enumerated ~100 potential AD targets ^3–5^. However, most studies have not accounted for possible latent factors which could indeed affect the biological discoveries as has been shown before by Yi et. al. ^6^. With the advances of high-throughput technologies, measuring the multiple omics data at the single-cell level has explained part of the heterogeneity of AD from the perspective of cell-type differences^7^ but the number of single-cell level studies for AD are still small.

Using an unsupervised learning algorithm; the Data-Adaptive Shrinkage and Clustering (DASC) algorithm ^6^, as well as expression profiles of human post-mortem samples, we identified a latent factor that reflects cell type heterogeneity. This latent factor was validated in multiple independent studies and was found to exist in AD, Mild Cognitive Impairment (MCI), and normal control samples as well. The discovery and validation of this latent factor could potentially explain the failure of some of the recent clinical AD trials. Here, we aim to highlight the existence of this factor and the need to account for it in identifying accurate AD biomarkers.

## Results

### A latent factor is identified and validated using unsupervised learning in brain postmortem samples

To identify latent factors in postmortem brain samples, we applied the DASC algorithm ^6^ to AD samples from the AMP-AD consortium. DASC uses a data-adaptive shrinkage method to cluster samples based on their biological characteristics and then uses semi-nonnegative matrix factorization (semi-NMF) to identify latent factors. We applied DASC to AD samples only (68 dorsolateral prefrontal cortex (DLPFC) samples) from the Memory and Aging Project (MAP) (Materials and Methods). We used only AD samples to create a homogenous discovery data set and to reduce the effect of molecular changes between AD and normal samples. The top 3000 most variable genes based on Median Absolute Deviation (MAD) were used as input to DASC, with the number of clusters ranging from k=2 to k=8 and with 500 iterations. We examined the robustness of the clusters using the cophenetic and silhouette coefficients, which yielded k=3 as the most robust number of clusters (Fig. 1A and Fig. S1A). Cluster 1 had 15 samples, cluster 2 had 32 samples, and cluster 3 had 21 samples.

**Fig. 1.**
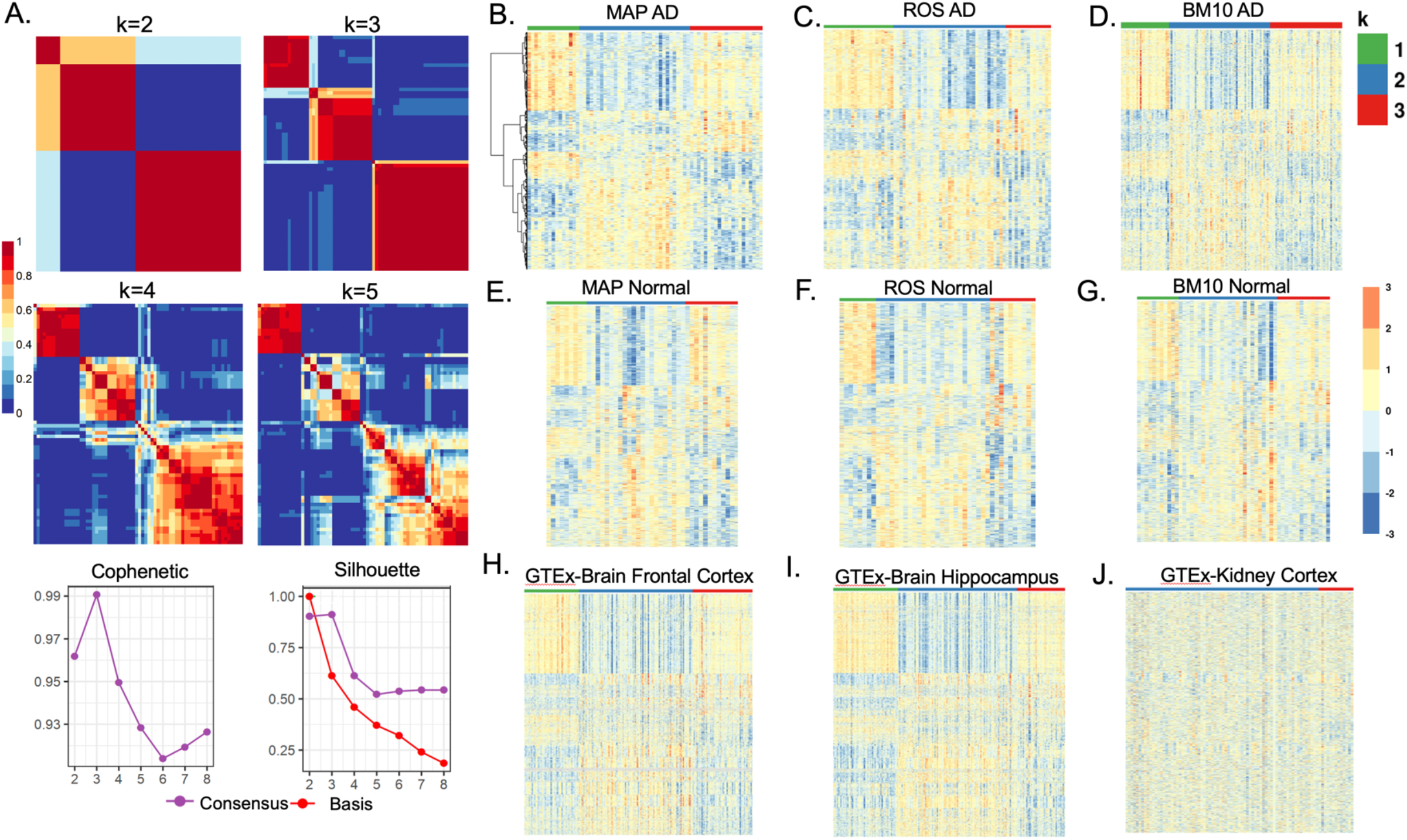
Identification and validation of a latent factor in AD postmortem samples using DASC. (A) The top 3000 most variable genes across 68 AD samples from the Memory and Aging Project (MAP) were used as input to DASC. We ran DASC with clusters ranging from k=2 to k=8 and found k=3 to produce the most robust clusters based on the cophenetic and silhouette coefficients. Cluster 1 had 15 samples, cluster 2 had 32 samples, and cluster 3 had 21 samples (B) The AD gene signature generated using the MAP AD sample. (C)-(J) The gene signature was validated in AD and normal samples in other cohorts. The genes in the validation studies had the same order as in the MAP AD dataset. This strongly validates the identified gene signature and the existence of the latent factor in postmortem brain regions regardless if they are AD or normal.

To validate these clusters and to verify that they were reproducible in other cohorts, we created a gene signature (gene expression profile) associated with them. We did so by identifying each cluster’s differentially expressed genes (DEGs), comparing each in a pairwise manner using DESeq2 ^8^ (Materials and Methods). We then selected the top DEGs which passed a false discovery rate (FDR) of 1% and which had an absolute fold change ≥1.5. This method yielded a gene signature of 2252 genes, which we used to build a random forest (RF) classifier on which the MAP discovery dataset was used as the training data. We then used the RF classifier to predict a cluster label for each sample in the validation studies as explained below.

We first examined the similarity of the gene signature in two validation studies: the Religious Orders Study (ROS), which has 75 AD DLPFC samples, and the Mount Sinai Brain Bank (MSBB) BM10 brain region study, which has 129 AD samples. We observed similar gene expression patterns in the MAP study and the two validation studies (Fig. 1B-D). The gene signature was also validated in the normal samples in the two validation studies as shown in (Fig. 1E-G). These findings strongly indicate that the latent factor is not limited to a single study, but rather exists in multiple studies.

We next validated the gene signature in three brain tissues from the MSBB study (BM22, BM44, BM36), in two brain tissues from the MayoRNAseq project [Ref], and in MCI samples, again finding a similar gene expression pattern (fig. S2). The replication of the gene signature in normal and MCI samples indicates that the latent factor exists in all postmortem brain samples and is not due to molecular changes between AD and normal or MCI samples.

As an additional validation step, we examined the gene signature using RNA-seq data from the Genotype-Tissue Expression (GTEx) project [Ref] for the following tissues: brain frontal cortex, brain substantia nigra, brain hippocampus, brain cerebellum, kidney cortex, lung, liver, and whole blood (Fig. 1H-J and Fig. S3). The MAP AD gene signature was only replicated in the brain tissue, indicating that the latent factor is intrinsic to brain tissue regardless of phenotype (AD, MCI, or normal) and regardless of age (the GTEx data is mainly derived from a young population, while AMP-AD samples are drawn from an old population.)

### Biological impact of the latent factor and its associations with clinical variables

To better understand the biological impact of the three sample clusters identified by the latent factor, we performed Gene Set Enrichment Analysis (GSEA) [Ref] on each cluster by finding genes that were differentially expressed in one cluster versus the other two clusters, as shown in Fig. S4.

Clusters 1 and 2 had opposite enrichment for the same GO and KEGG terms, indicating the unique molecular behavior in each. Cluster 1 was mainly enriched in synaptic and neuronal pathways, such as the synaptic membrane (GO:0097060), regulation of synaptic plasticity (GO: 0048167), neuron projection terminus (GO: 0044306), neurotransmitter secretion (GO: 0007269), and neuroactive ligand-receptor interaction (KEGG:hsa04080). Cluster 2 was enriched in extracellular organization (GO: 0030198, GO: 0043062), growth factor binding (GO: 0019838), endothelium development (GO:0003158), complement and coagulation cascades (KEGG:hsa04610), notch signaling pathway (KEGG: hsa04330), and hematopoietic cell lineage (KEGG:hsa04640). Cluster 3 was mainly enriched in protein folding functions (GO:0061077 and GO: 0006458), the Toll like receptor signaling pathway (KEGG: hsa04620) and antigen processing and presentation (KEGG:hsa04612). The opposite direction of the enrichment of the GO and KEGG terms reflects each cluster’s unique pathways and the unique molecular characteristics.

We next investigated associations between the clinical traits, technical artifacts, and the clusters identified by the latent factor (Fig. 2). We only found a significant association between neuroticism (an indicator of proneness to psychological distress) and the latent factor. Interestingly, there was no significant associations between the latent factor and AD pathological hallmarks such as amyloid and neurofibrillary tangles. This finding indicates that the clustering of samples is not due to disease progression or differences in AD pathology between samples. Additionally, the lack of significant association between the RNA Integrity Number (RIN), the post-mortem interval (PMI), and age of death could explain how the discovered latent factor exists in AD, normal, and MCI brain samples and confirms it is not due to technical artifacts.

**Fig. 2.**
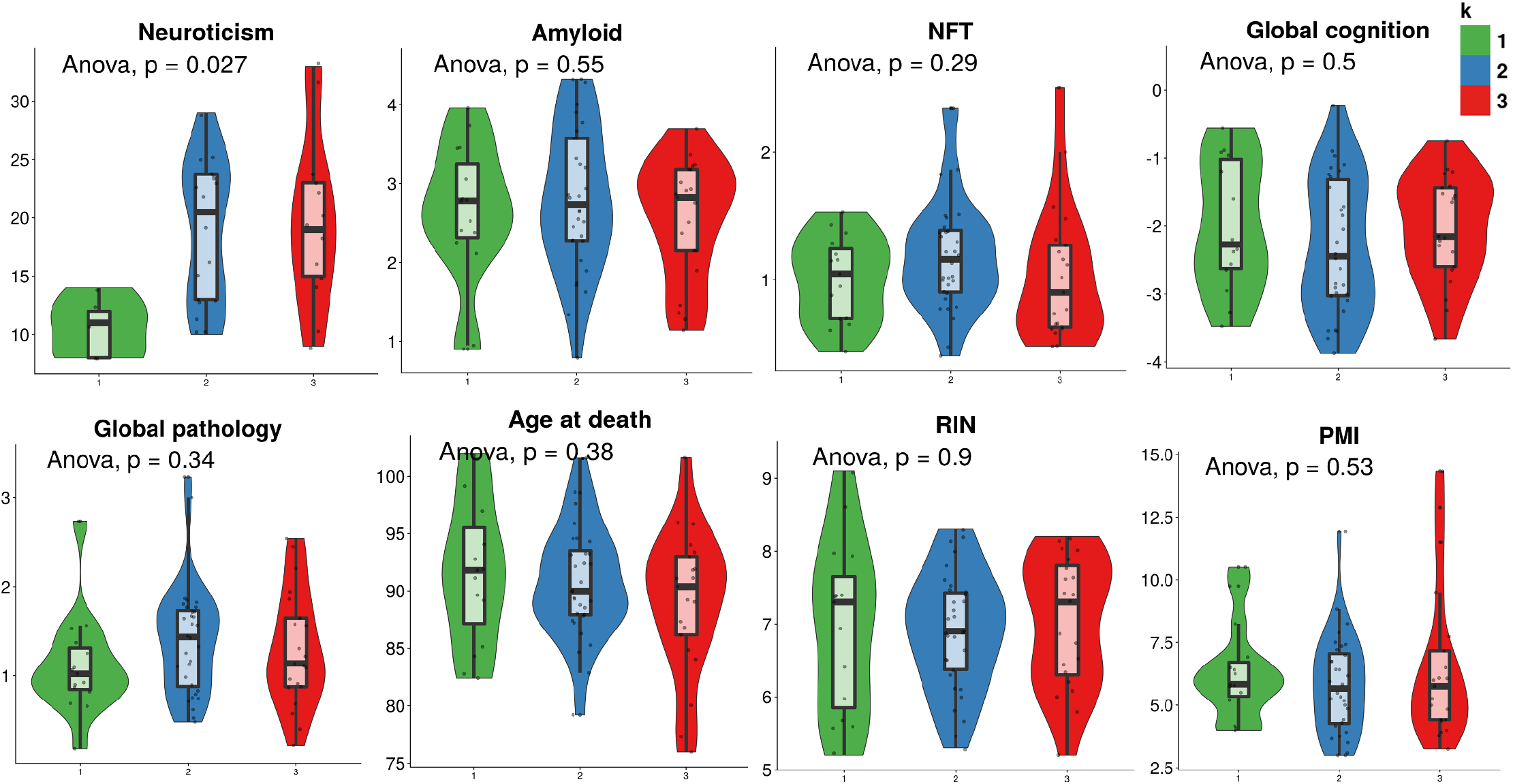
Association between clinical variables and the latent factor. Violin plots examining associations between the disocvered latent factor and clinical traits and technical artifacts. No significant associations were found between sample clusters and the clinical traits or technical artifacts excpet for neuroticism (indicator of proneness to psychological distress).

### The latent factor represents cell type proportion

To investigate whether the latent factor is due to cell-type proportions, we processed single-cell RNA-seq data for 48 samples from the ROSMAP (combined MAP and ROS) cohort ^9^ and examined the expression of the gene signature identified by the latent factor (Materials and Methods). We profiled 17,926 genes in 75,060 nuclei and found that 1,728 genes overlapped with the 2,252 MAP AD signature genes. We then used MAGIC (Markov affinity-based graph imputation of cells) ^10^ to impute the 1,728 signature genes to reduce the influence of missing values. Using the same ROSMAP bulk RNA-seq data used to identify the latent factor, we next trained a RF model to predict the cluster labels of the 48 single-cell RNA-seq samples. We then transformed the single-cell data to pseudo-bulk data per sample. Using this pseudo-bulk expression data, the trained RF assigned each sample a cluster label. We then calculated the celltype proportions for each sample (Materials and Methods) and compared the cell-type proportions among the three predicted clusters, as shown in Fig. 3A. Across the three clusters, both excitatory neuron cells and oligodendrocyte cells exhibited significant differences in cell proportions in AD samples (p-value=5.3e-05 and p-value=0.00048). The same cell-type proportion difference was also exhibited in normal brain samples (p-value=1.3e-06 and p-value=0.00025), which supports the effect of the latent factor regardless of disease state.

**Fig. 3.**
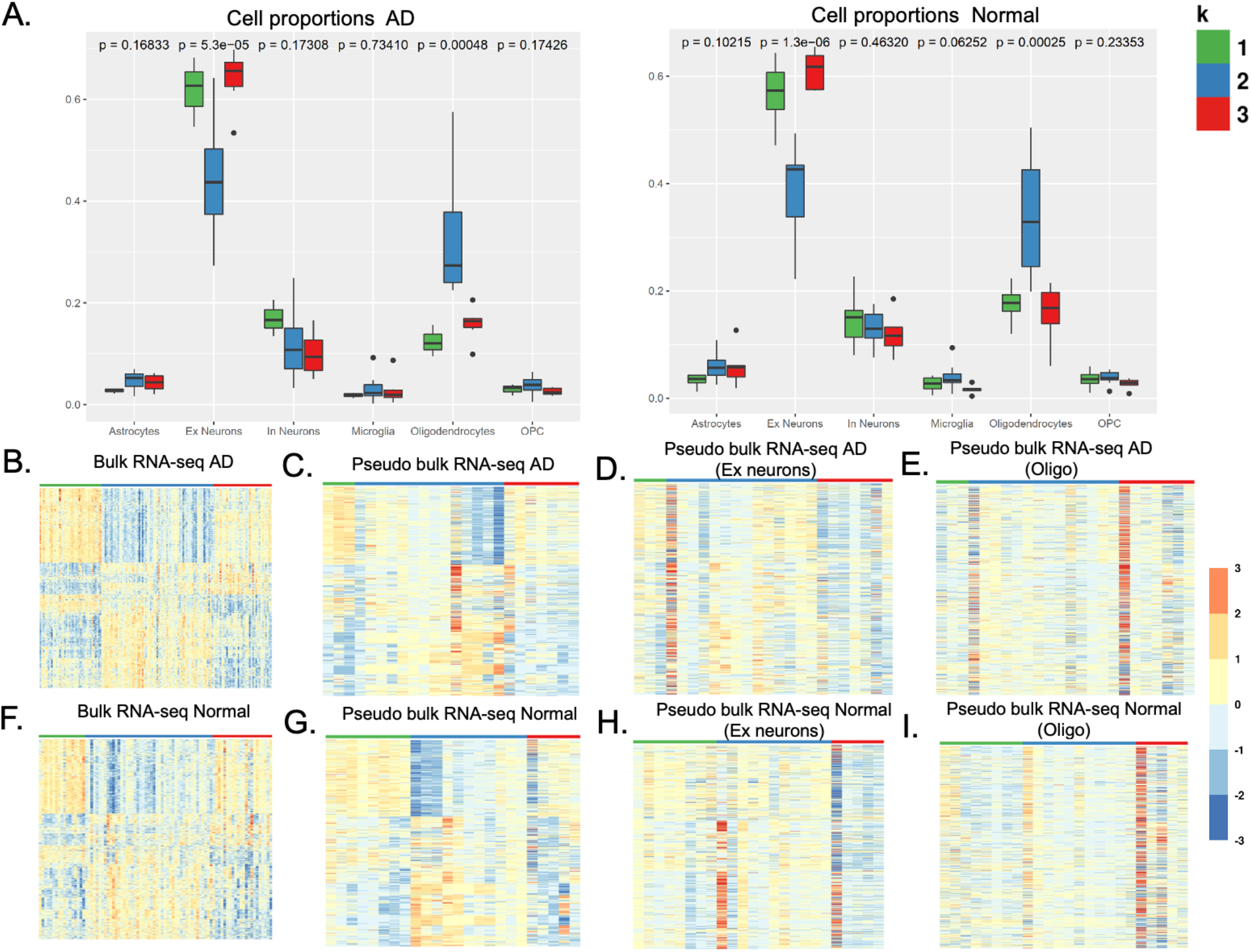
The discovered latent factor is assoicated with cell type proportions. Using single-cell RNA-seq data for 48 samples from the ROSMAP cohort [Ref], we comapred bulk RNA-seq and single cell psuedo bulk to examine the association between the latent factor and cell type proportions. (A) Cell type proproitons in AD and normal samples. Exhibitory (Ex) neurons and Oligodendrocytes show signicant association with the latent factor. (B)-(I) Gene signature validation in Psuedo bulk RNA-seq data where the gene signature is replicated in AD and normal pseudo bulk RNA-seq data as shown in C. and G. The gene signature is not replicated when examining on cell type at a time as shown in D,E,H, and I.

The observed difference of cell proportions across the three clusters identified by the latent factor explains its source. Additionally, the association of the latent factor with cell-type proportions explains why the latent factor existed in both AD and normal brain samples, and only in brain tissue versus other tissues.

We next compared the AD gene signature between bulk and pseudo-bulk RNA-seq data and found that the same signature was replicated (Fig. 3B and C). We then examined the signature only in the excitatory neurons and oligodendrocyte cells and found the gene signature was not replicated (Fig. 3D and E). The same gene signature was replicated in normal samples, as shown in Fig. 3F-I. These results further support that cell-type proportions are the source of the latent factor.

### New biological pathways revealed after adjusting for the latent factor

To further investigate the effect of the latent factor, we performed three types of AD versus normal DEG analysis: DEG analysis while adjusting for the latent factor, DEG analysis without adjusting for the latent factor, and DEG analysis with correction for cell-type proportions using a regression model (Materials and Methods). We combined the MAP and ROS samples (143 AD samples and 86 normal samples) to increase the statistical power of the DEGs. Fig. 4 D-E shows a Venn diagram with the number of common and new DEGs found. Without adjusting for the latent factor, 358 DEGs were found (259 upregulated and 99 downregulated). Adjusting for the latent factor, 735 DEGs were found (489 upregulated and 246 downregulated), where all the DEGs reported without correction were included in the DEG list, in addition to 377 new DEGs. DEG analysis while regressing cell-type proportions produced 84 upregulated genes and 69 downregulated genes. These results support that adjusting for the latent factor results in more accurate identification of DEGs and also resulted in the identification of new enriched pathways, as shown in Fig. 4 A-B and Fig. S5. We also examined the DEGs’ direction of change across studies after adjusting for the latent factor and found they changed in the same direction across studies (Fig. S6).

**Fig. 4.**
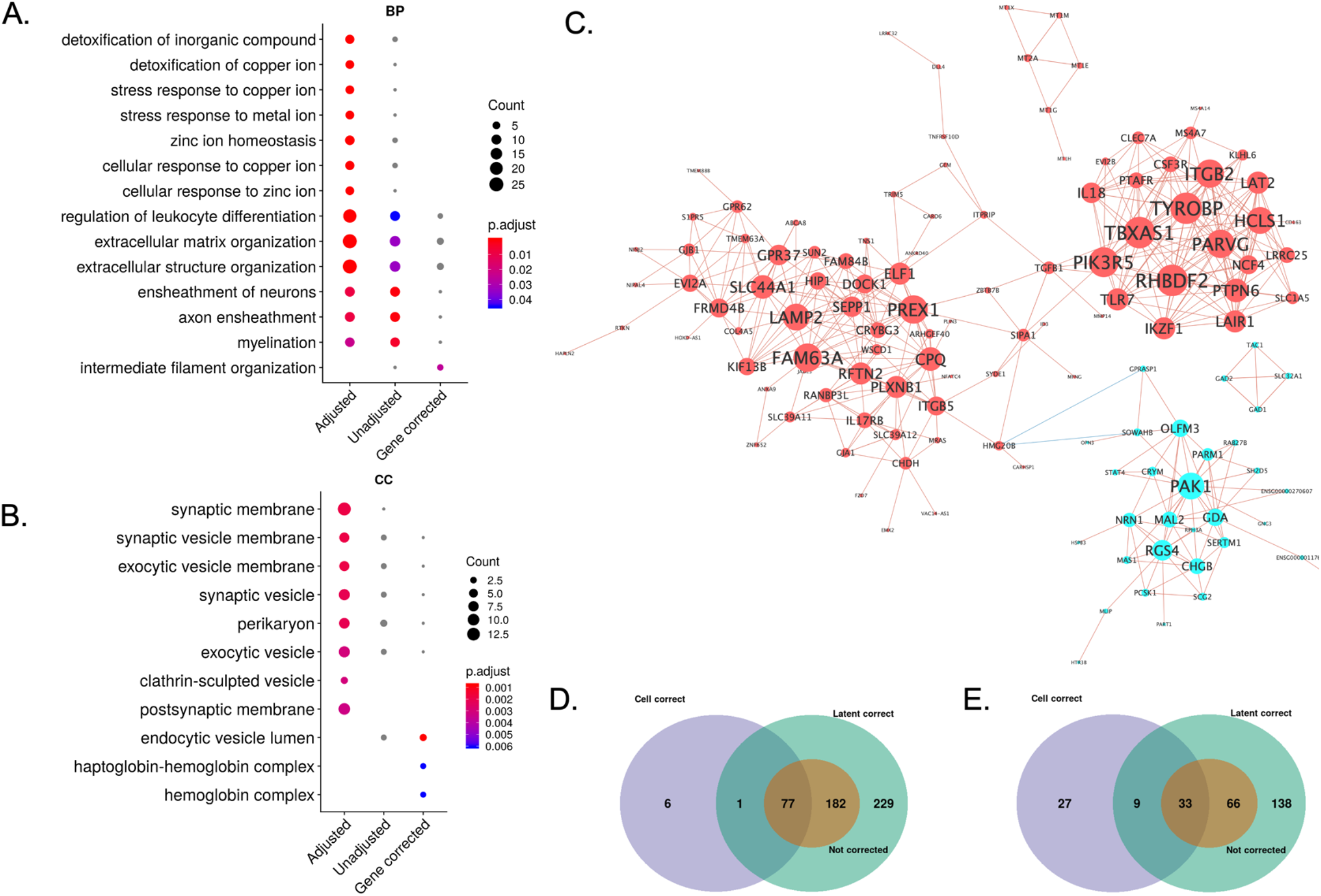
New biological pathways are identified after adjusting for the latent factor. (A) Comaprison of top enriched GO terms (BP) using upregulated DEGs found after adjusting for the latetnt factor. Adjusted refers to AD vs. Normal DEGs found after adjusting for the latent factor. Unadjusted refers to AD vs. Normal DEGs found without adjusting for the latent factor. Gene corrected regers to AD vs. Normal DEGs found after adjusting for cell type proprtions using a regression model. (B) Comaprison of top enriched GO terms (CC) using downregulated DEGs. (C) Gene co-expression network using 367 DEGs found only after adjusting for the latent factor. Red nodes are genes up-regualted and cyan nodes are downregulated genes. The color of edge reflects whether the correlation is positiove or negtive. The size of each node reflects the node degree. (D) Venn diagram showing the overlap between the set of upregualted DEGs found after adjusting for the latent factor, non-adjusted, and gene corrected. (E) Venn diagram showing the overlap between the set of downregualted DEGs found after adjusting for the latent factor, non-adjusted, and gene corrected.

When adjusting for the latent factor, upregulated genes were enriched in GO terms, such as detoxification of inorganic compounds (GO:0061687), detoxification of copper ions (GO:0010273), stress response to copper ions (GO:1990169), stress response to metal ions (GO:0097501), zinc ion homeostasis (GO:0055069), and cellular response to zinc ions (GO:0071294). Both copper and zinc have been found to affect the aggregation of amyloid-B and the oxidative stress process ^11–13^.Specifically, it has been reported that copper imbalance and dysregulation can lead to cognitive impairment ^14^. Additionally, copper has been suggested as a biomarker for AD, in which AD patients are stratified based on copper imbalance^15^. Abnormalities in zinc levels have been related to neurotoxicity as well^11^.

Additional enriched GO terms included the B-cell receptor signaling pathway (GO:0050853), regulation of B-cell activation (GO:0050864), B-cell activation (GO:0042113), and NK T cell activation (GO:0051132), all of which are related to immune response. Associations or causation between B-cells and NK T and AD have not yet been established, and further investigation is warranted ^16–20^.

Downregulated genes were enriched in GO terms, including the neuropeptide signaling pathway (GO:0007218), neurotransmitter transport (GO:0006836), synaptic membrane (GO:0097060), inhibitory synapse (GO:0060077), and perikaryon (GO:0043204). Our identification of new pathways after correcting for the latent factor highlights the importance of adjusting for the latent factor before downstream analysis, allowing for more accurate identification of AD biomarkers.

To investigate whether the DEGs found after adjusting for the latent factor are cell-type-specific, we examined the overlap between DEGs found using bulk RNA-seq and those found using single-cell RNA-seq. We compared the 1,031 AD versus normal DEGs reported in the AD single-cell RNA-seq study ^9^ to the DEGs we found after adjusting for the latent factor. Among the 489 upregulated genes found after adjustment, only 28 overlapped with the 1,031 cell-type-specific DEGs. Among 246 downregulated genes found after adjustment, only 19 genes overlapped. These findings further support that DEGs found after adjusting for the latent factor are not cell-type-specific and are not reflective of cell-type proportion.

We next focused on the 367 DEGs found after adjusting for the latent factor and created a gene co-expression network, as shown in Fig. 4C (Materials and Methods). The genes were coexpressed in three clusters, two of which contained upregulated genes and one of which contained downregulated genes.

The first cluster contained the following hub genes: TYROBP, TBXAS1, PRVG, RHBDF2, PIK3R5, ITGB2, and LAT2. TYROBP and ITGB2 are both involved in immune and microglia networks in AD ^4,21^ Previous studies have found RHBDF2 to be differentially methylated and associated with AD pathology ^22,23^.

The second cluster contained the following hub genes: FAM63A, LAMP2, PREX1, SLC44A1, DOCK1, CPQ, SEPP1. FAM63A was previously found to be nominally significant in AD GWAS studies and significantly associated with AD disease duration ^24–26^. LAMP2 has been nominated as a cerebrospinal fluid AD biomarker^27,28^. PREX1 has been associated with autism and synaptic plasticity, as reported in ^29,30^.

The third cluster contained the following hub genes: PAK1, RGS4, OLFM3, GDA, MAL2. PAK1 has been shown to be dysregulated in multiple neurodegenerative and neurodevelopmental disorders, including AD, HD, autism, and X-linked mental retardation ^31–34^. Previous studies also found that RGS4 has lower expression levels in AD samples ^35,36^.

### The latent factor impacts previously discovered AD modules

To investigate the impact of the latent factor on previously defined AD modules, we compared the distribution of the intra-module gene correlations within each cluster of samples predicted by the latent factor. Specifically, we calculated the pairwise absolute Pearson correlation coefficients (PCC) of the m109 module genes derived from the ROSMAP bulk RNA-seq data ^3^, and the beige module derived from the MSBB BM22 brain region microarray data ^4^, as shown in Fig. 5. The m109 module is a co-expression gene module consisting of 390 genes, which was found to be associated with cognitive decline. The beige module is also a coexpression module, consisting of 95 genes, which was found to be associated with AD pathology. Its top functional category consisted of glucose homeostasis.

**Fig. 5.**
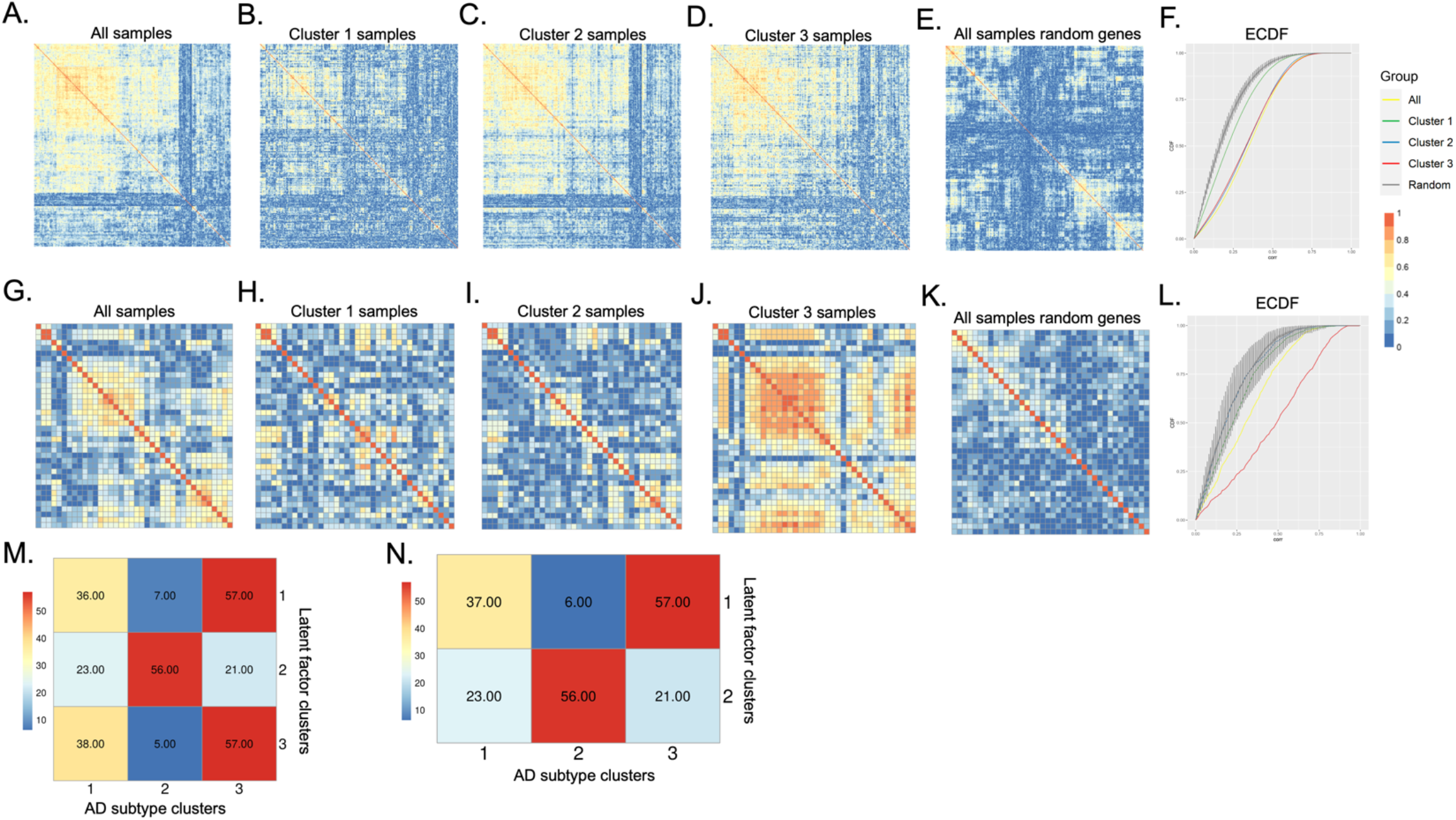
Impact of latent factor on previous studies. (A-D) Patterns of pairwise Pearson Correlation Coefficient (PCC) of Mostafavi’s et al. [Ref] m109 module genes using all samples and stratified samples using the latent factor. (E) heatmap of PCCs of randomly selected genes using all samples. (F) Empirical Cumulative Density Function (ECDF) plots of PCCs in A to E. (G-J) Patterns of PCC of Zhang’s et al. [Ref] beige module genes using all samples and stratified samples using the latent factor in the MSBB-BM22 samples. (K) heatmap of PCCs of random selected genes using all MSBB-BM22 samples. (L) ECDF plots of PCCs in G to K. (M) Confusion matrix comparing the cluster assignmnets using the latent factor and AD subtype clusters reported in Neff et al. [Ref] where the AD subtype clusters were divided into atypical-A (1), typical-C (2), and intermediate-B (3). The cell values represent percenatges of samples common between the two studies. Latent factor cluster 2 has the most commnality with the typical AD subtype class. Clusters 1 and 3 had the most commanlity with the intermediate AD subtype class. (N) Similar to M but clusters 1 and 3 found using the latent factor were combined into cluster 1.

We observed that the m109 module genes are less correlated within Cluster 1 samples (Fig. 5B), as compared to Cluster 2 and 3 samples (Fig. 5C-D). By examining the PCCs’ empirical cumulative density function curve (ECDF), Cluster 1’s CDF is more similar to the CDF of a set of random genes, whereas Cluster 2 and 3’s CDFs are more similar to one another (Fig. 5F). This result illustrates how the samples stratified by the latent factor exhibit different co-expression patterns across the clusters and how samples within each cluster have intrinsic molecular characteristics unique to each cluster.

We observed similar behavior in the beige module. This module’s genes are more correlated in Cluster 3 (Fig. 5J) compared to Clusters 2 and 3 (Fig. 5H-I), and when examining the CDF curve, genes in Clusters 1 and 2 overlap with the CDF generated using a random set of selected genes (Fig. 5L).

We next examined the similarities between clusters identified by the latent factor and AD subtype clusters reported in Neff et al., which identified three subclasses and five subtypes using the BM36 brain region from the MSBB study ^37^. The number of samples common with BM36 samples we analyzed was 83 samples. The three AD subtype classes were labeled as: atypical AD (24 samples); intermediate AD (30 samples); and typical AD (29 samples). We then compared the commonality of samples in each AD subclass with the cluster samples identified using the latent factor, as shown in Fig. 5M-N

(Fisher’s exact test p-value=2.1×10-5). The latent factor Cluster 2 (48 samples) had 56% commonality with the typical AD subclass samples. The typical AD class was characterized by synaptic depression reflective of neuronal activity, which is similar to our findings in Fig. S4, in which Cluster 2’s synaptic pathways were not enriched. Moreover, as shown in Fig. 3A, Cluster 2 samples had lower cell proportions for excitatory neurons and inhibitory neurons. Latent factor Clusters 1 and 3 had the most commonality with the intermediate AD subclass samples. The intermediate AD class was characterized by synaptic excitation, as reported in Neff et al., which is similar to Cluster 1 and 2 samples, in which synaptic pathways were enriched (Fig. S4.) Although more than half of the latent factor clusters shared commonality with the AD subclasses, the remaining samples were divided between the AD subclasses 1 and 3. This finding could be attributed to differences in the cell proportions of oligodendrocytes. As shown in Fig. 3A, all three latent factor clusters had significant differences in oligodendrocyte cell proportions (p-value=0.00048).

### Impact of the latent factor on metabolic patterns

To investigate the downstream impact of the latent factor, we characterized metabolic patterns observed in the identified clusters. Out of all samples from the ROSMAP cohort included in this study, 103 were collected from individuals whose prefrontal cortex tissue were also profiled by untargeted metabolomics. Twenty-one of them were identified in cluster 1, 53 were found in cluster 2 and 29 in cluster 3. Significant metabolic difference was observed among these clusters, especially between cluster 1 and the other two clusters (Fig. 6). Compared to cluster 2 and 3, majority (28/32) of the significant differential metabolites (p-value < 0.05 and log 2(fold change) > 1) were less abundant in cluster 1 (Fig. 6 and 7, Supplementary Table 1). The four metabolites with higher level of concentrations include N-acetyl-3-methylhistidine, trigonelline, 3-methylhistidine, all of which have previously been reported to be related to low nutrition state of brain ^38–40^. In down-regulated metabolites, several were found to provide alternative oxidative fuels for cognitive functions, including glycogens such as maltotetraose, maltotriose^41^.

**Fig. 6.**
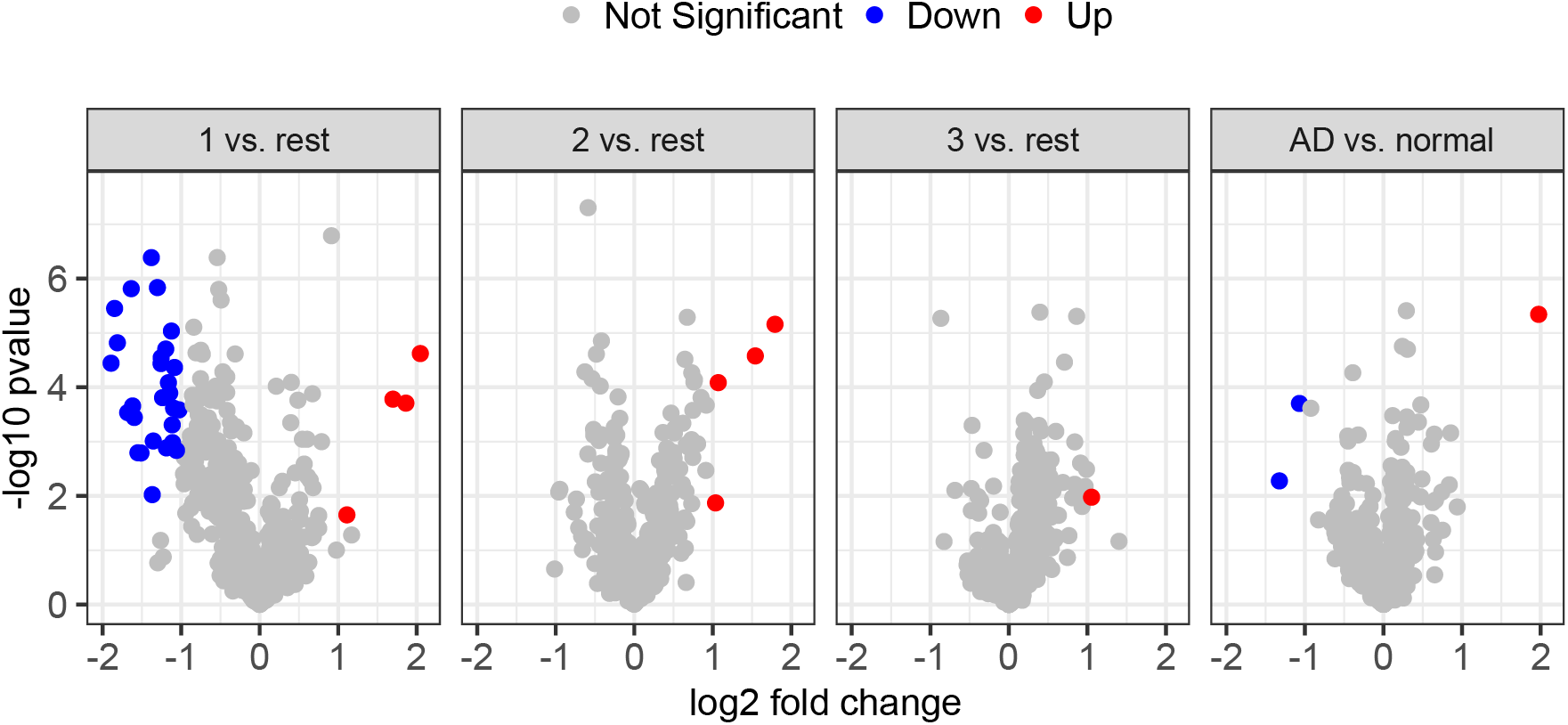
Volcano plot of differentially expressed genes identified in one-versus-rest comparison across 3 clusters. Differential metabolites significantly (p<0.05) up- or down-regulated by at least 2 folds in cluster 1 samples compared to the rest of the samples are colored in red and blue respectively.

**Fig. 7.**
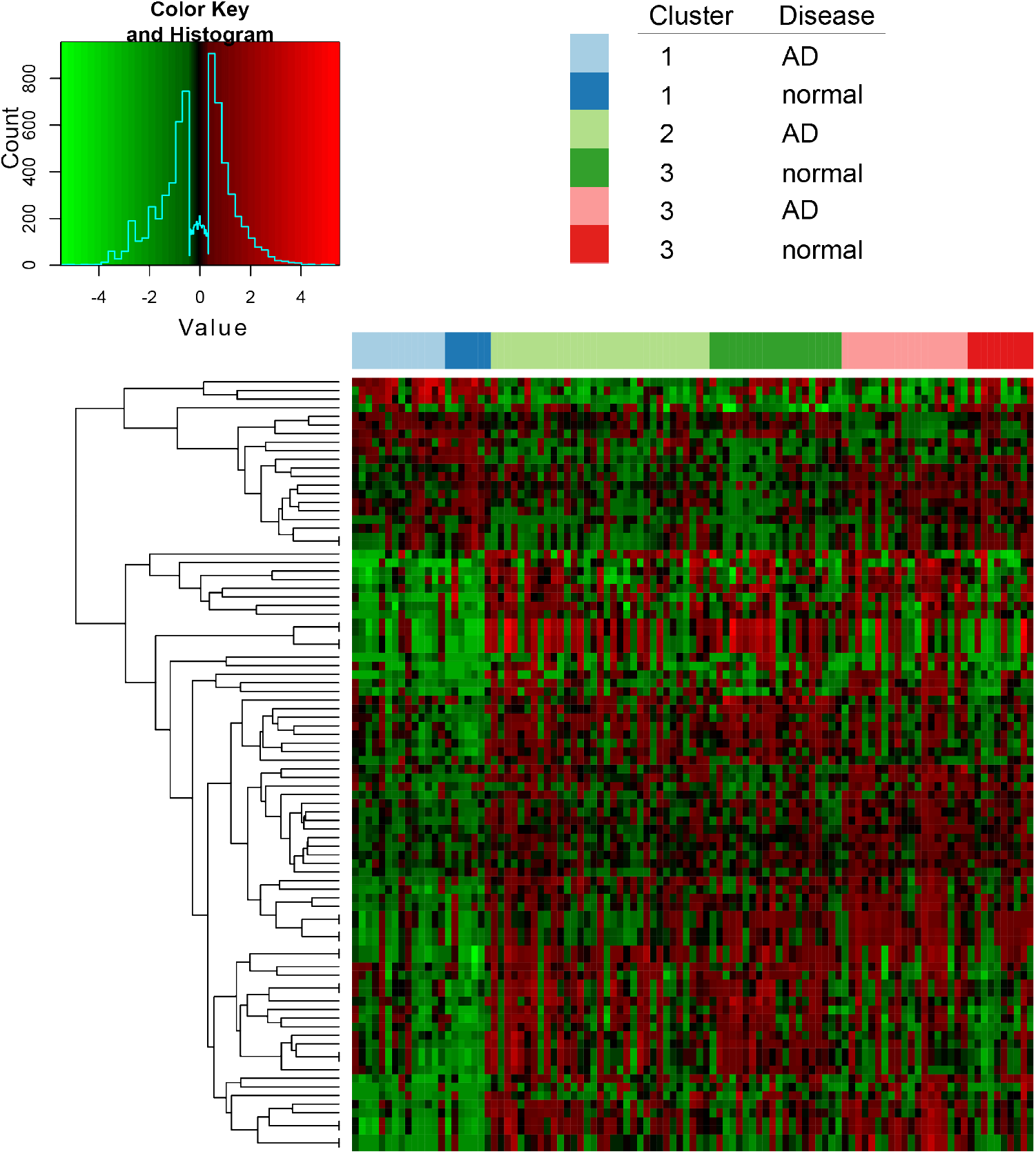
Heatmap of top 30 differential metabolites (rows) identified in each one-versus-rest comparisons across 3 clusters. Samples (columns) are grouped by the combination of cluster and AD diagnosis.

We next examined metabolic co-perturbations specific to cluster 1 via a partial correlation network learned from contrasting metabolic profiles of cluster 1 samples against the rest of the samples. Most of the differential metabolites with p-value smaller then 1e^-3^ were identified within the same sub-community (Fig. 8) by a fast greedy algorithm (Materials and Methods). The community included strongly connected metabolites including cortisol and several metabolites within kynurenine/tryptophan (Fig. 8). The production of kynurenine from tryptophan can be triggered by higher level of cortisol^42^ and is associated with various neuropsychiatric disorders such as depression-associated anxiety, psychosis and cognitive decline^43^. Lower levels of cortisol and downregulated kynurenine pathway were observed in cluster 1 samples, consistent with lower clinical variable neuroticism score, an indicator of proneness to negative moods, including anger, anxiety, emotional instability, and depression.

**Fig. 8.**
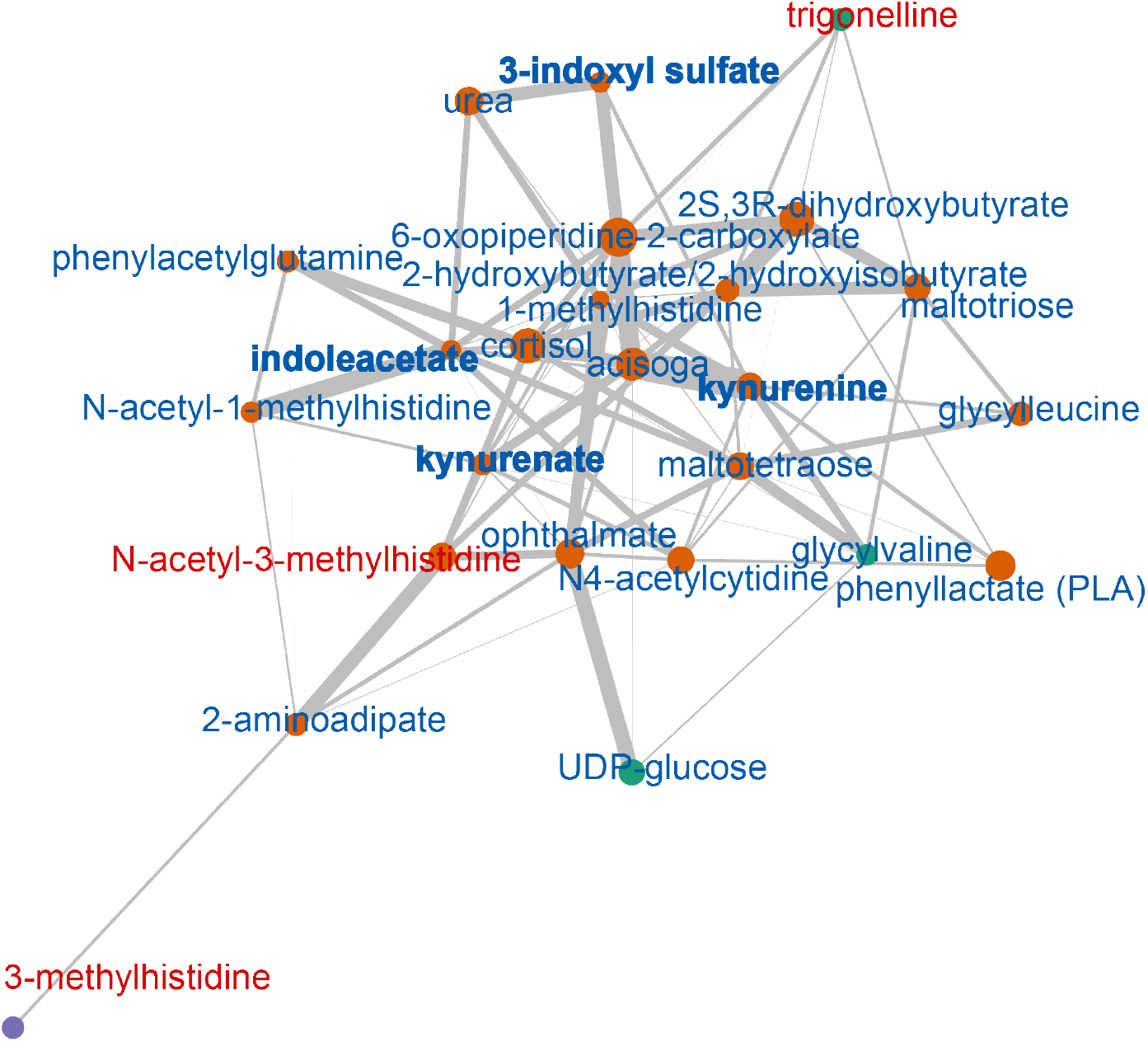
Modularity of cluster 1 differential metabolites (p<1e^-3^) in partial correlation network. Color of the nodes denotes node membership in fast greedy algorithm identified communities. Up- or down-regulation of metabolites abundance is indicated by color of node labels. Metabolites involved in tryptophan/kynurenine pathway are formatted in bold.

## Discussion

One of the most pressing challenges in finding new, improved AD therapies is identifying accurate biomarkers representative of the disease. Several consortia have deposited bulk RNA-seq data from postmortem human brain samples, and several studies using this data have reported possible new AD targets. One caveat in using this bulk RNA-seq data, however, is the possible hidden effect of latent factors.

In this study, we used an unsupervised machine learning algorithm (DASC) to identify just such a latent factor. Using AD samples, we identified three robust clusters associated with a particular latent factor, which we then validated across multiple studies using AD, MCI, and normal samples. We found the factor represented cell-type heterogeneity in which the excitatory neurons and oligodendrocyte cells exhibited different cell-type proportions across the three clusters it had identified. In the majority of AD studies, only reported artifacts are adjusted before downstream analysis to identify AD associated genes and biomarkers. Yet it is clearly imperative to first discover and then adjust for any latent factors before undertaking further analysis.

As we demonstrated here, adjusting for the latent factor led to more accurate downstream analysis, allowing us to identify several significant new biological pathways. Examples included the detoxification of copper ions and cellular response to copper and zinc ions, neither of which were found without adjusting for the latent factor.

The effect of the latent factor was not limited to transcriptomic data as supported by the significant metabolic differences observed among the three clusters identified. This supports how the latent factor had an effect multiple dimensions of omics’ data and is not random by nature.

Notably, a simple regression model that adjusted for cell-type heterogeneity using a set of gene markers performed poorly in identifying biological pathways. Moreover, using a regression model with a large number of gene markers as covariates did not totally eliminate the effect of cell-type heterogeneity. These findings merit further investigation to understand the limits in using regression models while adjusting for cell-type heterogeneity.

Ultimately, we were able to identify a unique set of 367 AD DEGs which were found only after adjusting for the latent factor. Based on a co-expression analysis, the DEGs clustered into three modules, one of which (TYROBP) is related to the immune pathways. We need to closely dissect these genes and validate them to learn more about their role in AD.

As single-cell RNA-seq analysis continues to improve along with the decrease in experiment costs, the issue of cell-type proportions will be of less concern, but hidden latent effects could still exist, especially when considering the large number of available samples and institutes working in the field. Thus, accounting for hidden factors remains a critical step. We anticipate that our proposed approach will help identify accurate AD biomarkers in other studies and after accounting for the latent factor, lead to an improved understanding of the disease.

## Materials and Methods

### Data sources

We used four studies from the Accelerating Medicines Partnership Program for Alzheimer’s Disease (AMP-AD) [Ref]. One study was used for the identification of the latent factor and three studies were used for validation. The studies included: the Rush Memory and Aging Project (MAP) [Ref] study, which consisted of 313 samples from the DLPFC brain region; the Religious Orders Study (ROS) [Ref], which consisted of 325 samples from the DLPFC brain region; and the Mount Sinai Brain Bank (MSBB) [Ref] study, which consisted of 1,026 samples across four brain regions: frontal pole (BM10, 265 samples), superior temporal gyrus (BM22, 264 samples), parahipocampal gyrus (BM36, 267 samples), and frontal cortex (BM44, 230 samples). We first used the BM10 brain region from the MSBB study for validation due to its proximity to the DLPFC region. We also used RNA-seq data from the cerebellum and temporal cortex brain tissues in the MayoRNAseq study [Ref]. Samples were filtered such that AD samples were used for the discovery and validation of the latent factor. The MAP and ROS AD samples were selected according to the following criteria: clinical consensus diagnosis (cogdx)=4, Braak stage ≥4, the Consortium to Establish a Registry for AD (CERAD) score <= 2. The control samples were selected such that cogdx=1, Braak stage <= 3, and CERAD >= 3. For the MSBB study, AD samples were selected according to the following criteria: clinical dementia rating (CDR) >= 1, neuropathology category as measured by CERAD (NP.1) ≥ 2, and Braak score ≥4. The control samples were selected such that CDR<=0.5, NP.1<=1, and Braak score <=3. For the MayRNAseq study, we used the diagnosis provided by the study. The number of AD and control samples in each study is shown in Table S1. Moreover, we used MCI samples as validation from three studies: MAP (n=95), ROS (n=75), and MSBB BM10 (n=39)

We also used RNA-seq data from the GTEx project [Ref] for the following tissues: brain frontal cortex BA9 (n=209), brain substantia nigra (n=139), brain hippocampus (n=197), brain cerebellum (n=241), kidney cortex (n=85), lung (n=578), liver (n=226), and whole blood (n=755). Data was normalized using DESeq2^8^ and batch corrected using limma^44^. The same RF model built using the MAP AD data was used to predict sample labels for the GTEx samples.

### Data filtering

Genes were filtered by removing the Y chromosome genes to reduce the sex effect and by removing transcripts with low counts to obtain 17,538 genes. We normalized the gene counts using DESeq2^8^ and applied log2 transformation to the counts. We corrected for the reported batch factors in each study using the limma package function ‘removeBatchEffect’^44^ in which we corrected for batch and adjusted for the RIN and PMI as covariates.

### Data-Adaptive Shrinkage and Clustering (DASC) algorithm

DASC^6^ starts by obtaining the batch-free matrix U (biological signal) using a non-parametric data-adaptive shrinkage method. The batch-free matrix is then used to obtain the batch-matrix B. Semi-NMF is then applied to the batch matrix B to find hidden batch factors.

### Gene signature

We created a gene signature (gene expression profile) associated with the latent factor by identifying DEGs by comparing each cluster found using DASC with k=3 in a pairwise manner and selecting the top DEGs which passed an FDR of 1% and had an absolute fold change ≥1.5. This resulted in a gene signature of 2,252 genes which we used to build a random forest (RF) classifier in which the MAP discovery dataset was used as the training data. The RF classifier was then used to predict a cluster label for each sample in the validation studies. When plotting the heatmap for the validation studies, we used the same gene order observed from the MAP AD study.

### Single-cell RNA-seq analysis

Single-cell RNA-seq data from 48 samples from the ROSMAP cohort ^9^ was processed using CellRanger software v2.0.2 (10x Genomics). We filtered out cells with fewer than 200 detected genes and proportion of counts mapping to mitochondria genes higher than 38.08%. Mitochondrially encoded genes and genes detected in fewer than two cells were excluded. This resulted in 17,926 genes profiled in 75,060 nuclei. All 75,060 cells were combined into a single dataset. Normalization and clustering were performed with Seurat v3^45^. Counts for all nuclei were scaled by the total library size multiplied by 10,000 and transformed to log space. A total of 3,000 top highly variable genes were identified using Seurat to perform principal component analysis (PCA). The top 10 PCs were used to build the k-nearest-neighbors (KNN) cell-cell graph with k=20. Then the shared nearest neighbor was built based on the KNN graph, in which clusters of the cells are identified. The initial clustering resulted in 13 clusters. To annotate these clusters, we used cell-type markers reported in^46^ to determine the cell type of each cluster. We ended up with six main cell types, including excitatory neurons (49.9%), inhibitory neurons (12.4%), astrocytes (4.8%), microglia (2.7%), oligodendrocytes (25.8%), and oligodendrocyte progenitor cells (3.6%).

### DEG analysis while adjusting for the latent factor

To compare the DEGs before and after adjusting for the hidden batch factor, we used the limma function “removeBatchEffect”^44^ such that the three identified clusters were treated as a batch. We report the DEGs before and after batch correction. For the gene-corrected results, we used marker genes and the regression model (BRETIGEA R package) from ^47^ to correct for celltype proportions. We used the R package clusterProfiler ^48^ to perform GO analysis.

### Gene co-expression network

Using the 367 DEGs found after adjusting for the latent factor, we calculated the correlation values between all DEGs using the R function cor.test. We filtered out gene pairs with correlation values < 0.7 (either direction) and correlation test p-value > 1×10-5 by assigning a correlation value of zero to pairs which do not pass these two thresholds. We then created a graph object using the R package igraph (available at https://igraph.org/r/) and then used Cytoscape^49^ to visualize the gene co-expression network.

### Metabolomics data process and differential analysis

The dorsolateral prefrontal cortex (DLPFC) metabolomic data from the ROSMAP studies were downloaded from Synapse (syn26007830). Raw intensities of named metabolites detected and measured in more than 60% of the included samples were log2 transformed and used for downstream analysis. Metabolites with two-sided t test P<0.05 and log2FoldChange >1 were considered as significantly differential metabolites.

### Metabolite co-perturbation network

We used the Graphical Lasso algorithm implemented in the R package huge (v1.3.5) to compute edge weights, which indicate the strength of the positive or negative partial correlation between metabolites. The network model was learned to represent the metabolic co-perturbation patterns specific to cluster 1 in two steps: First, two types of graphical models were constructed, one containing all included samples (case + control network) and a second from only cluster 2 and cluster 3 samples (control network). Next, the edges found in the case + control network that were also found in the control network were pruned. Fast greedy modularity optimization algorithm implemented in R package igraph (v 1.3.0) was then used for finding community structure within the learned metabolite co-perturbation network.

---

**Fig. S1.**
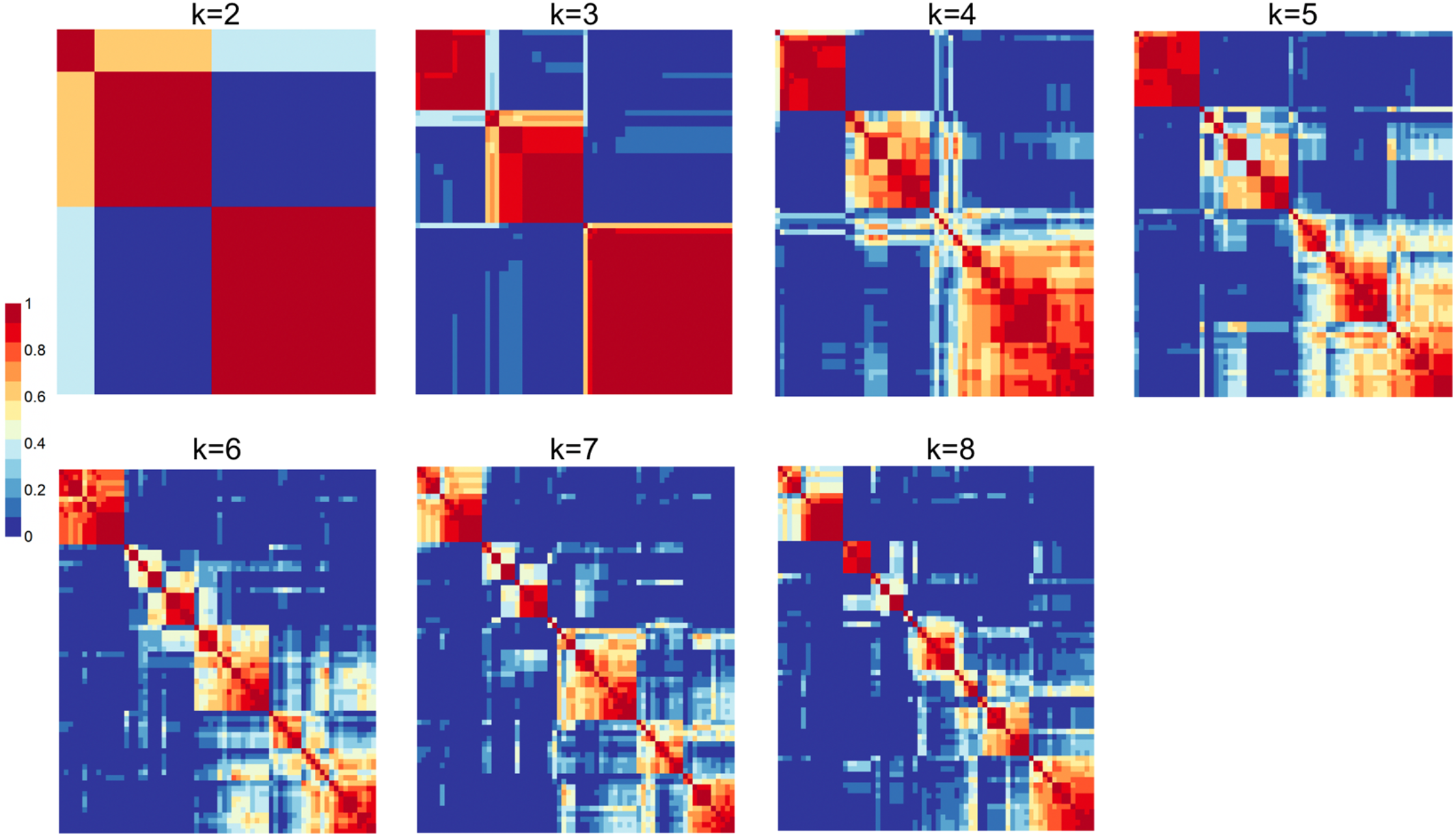
Clusters identified using DASC for k=2 to k=8.

**Fig. S2.**
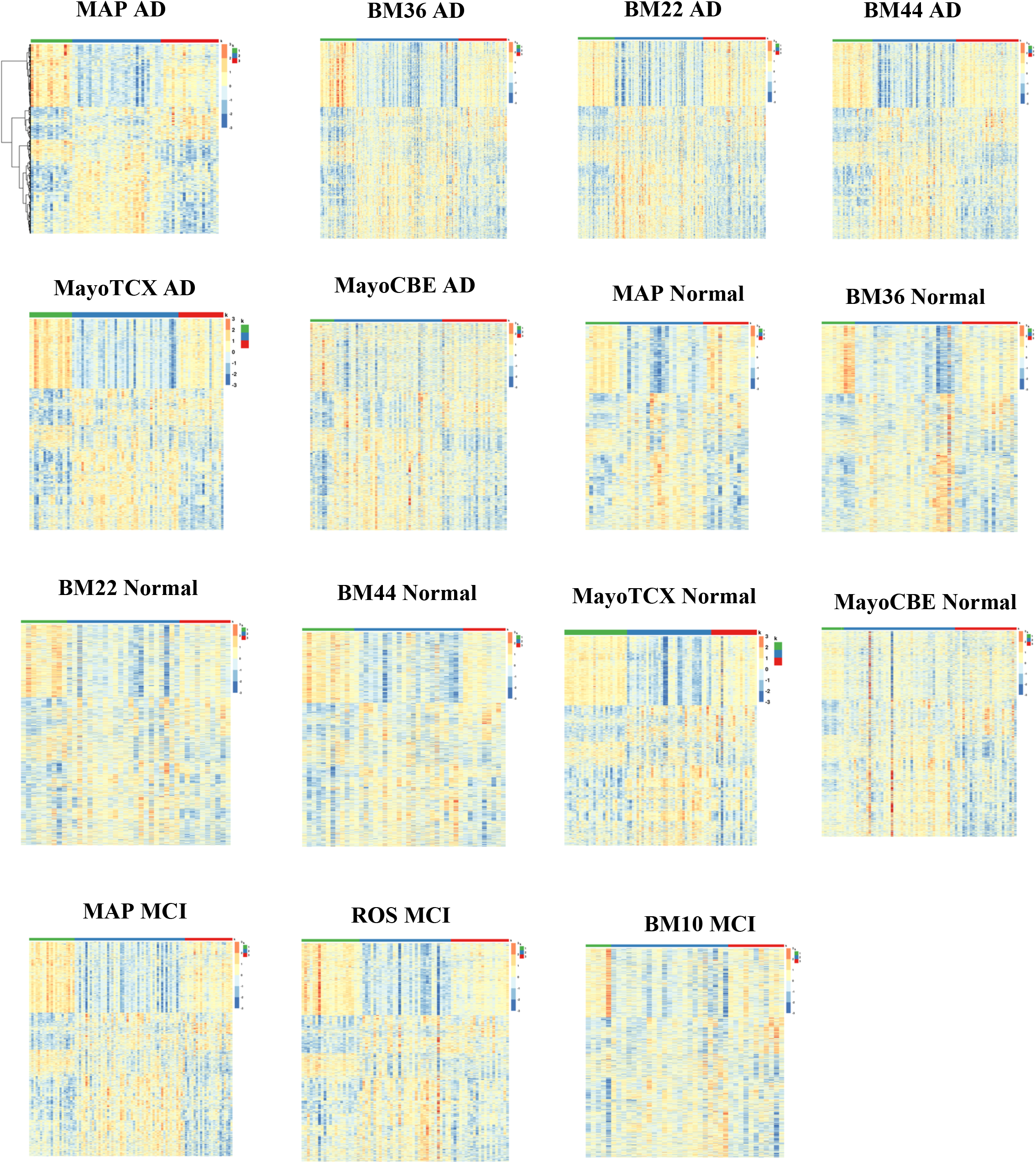
AD gene signature comparison across three studies. The MAP AD gene signature was generated using pair wise gene DEG analysis.

**Fig. S3.**
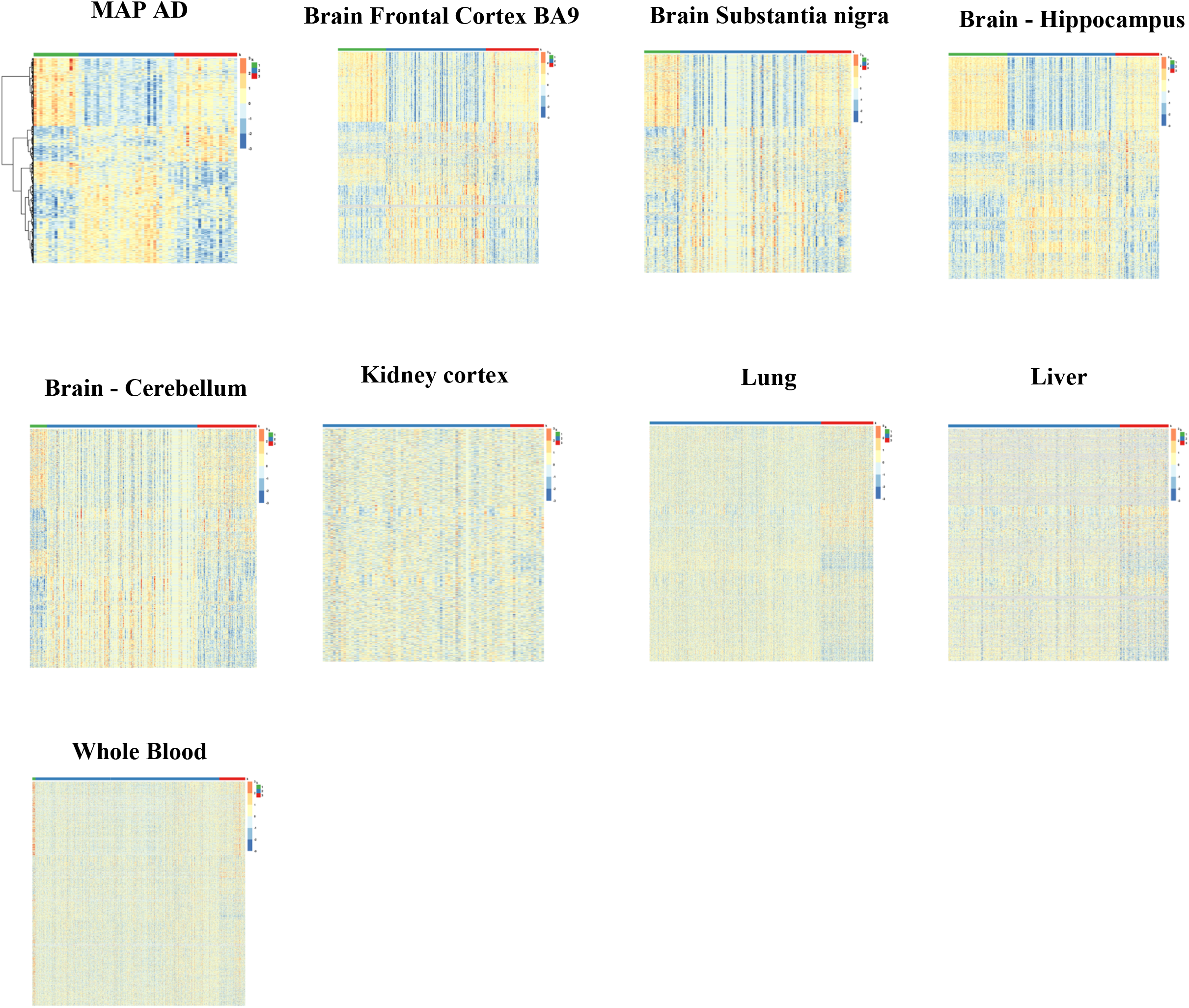
Validation using GTEx data across multiple tissues. The genes are ordered in the same order as the MAP AD gene signature.

**Fig. S4.**
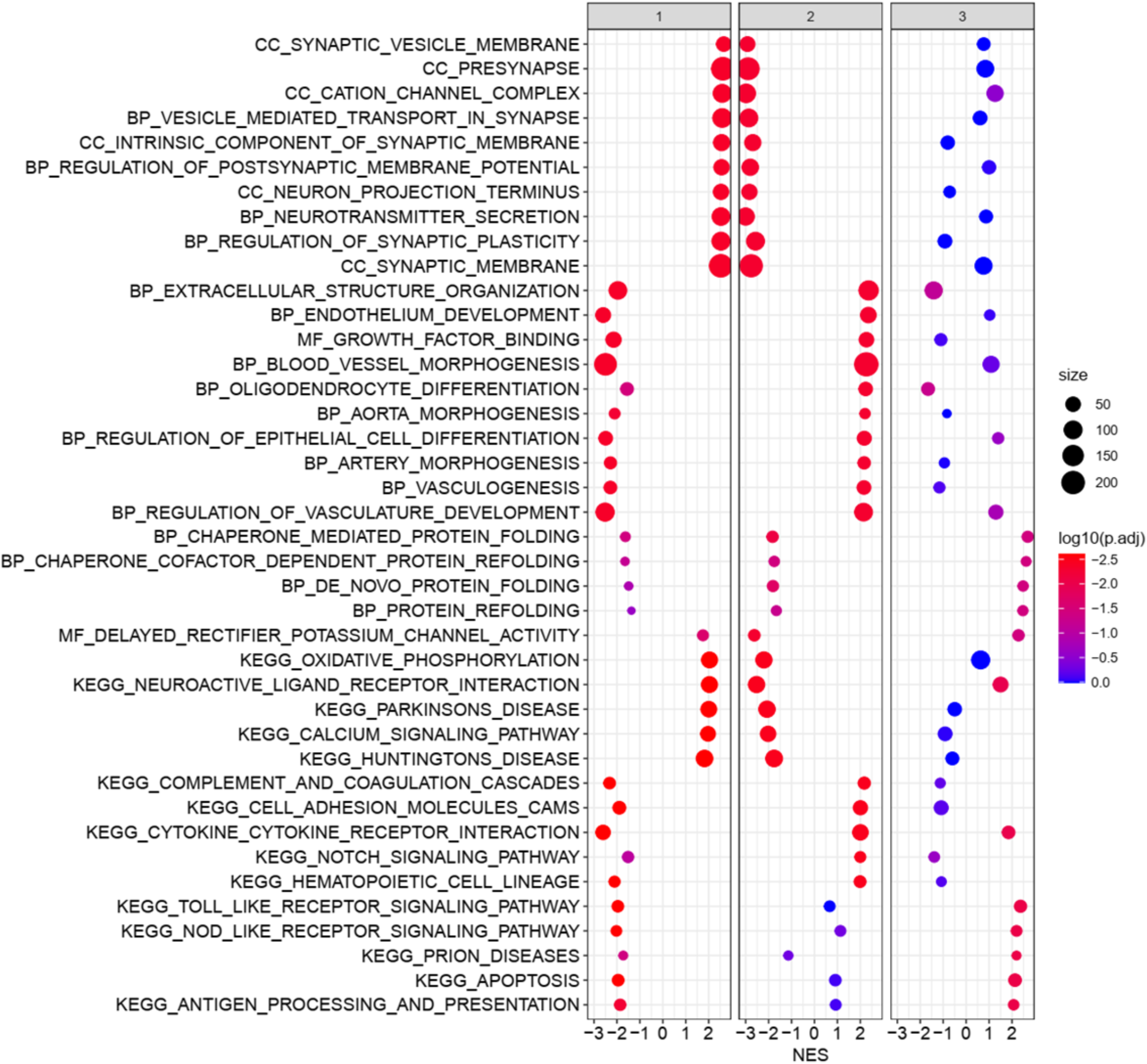
Functional annotation of latent variable DEGs (GO and KEGG) using GSEA. Each column represents one cluster. Clusters 1 and 2 have opposite enrichment for the same GO and KEGG terms which indicates the unique molecular behavior of the two clusters. Cluster 1 is mainly enriched in synaptic and neuronal pathways. Cluster 2 is enriched in extracellular organization, growth factor binding, and complement and coagulation cascades. NES refers to Normalized Enrichment Score.

**Fig. S5.**
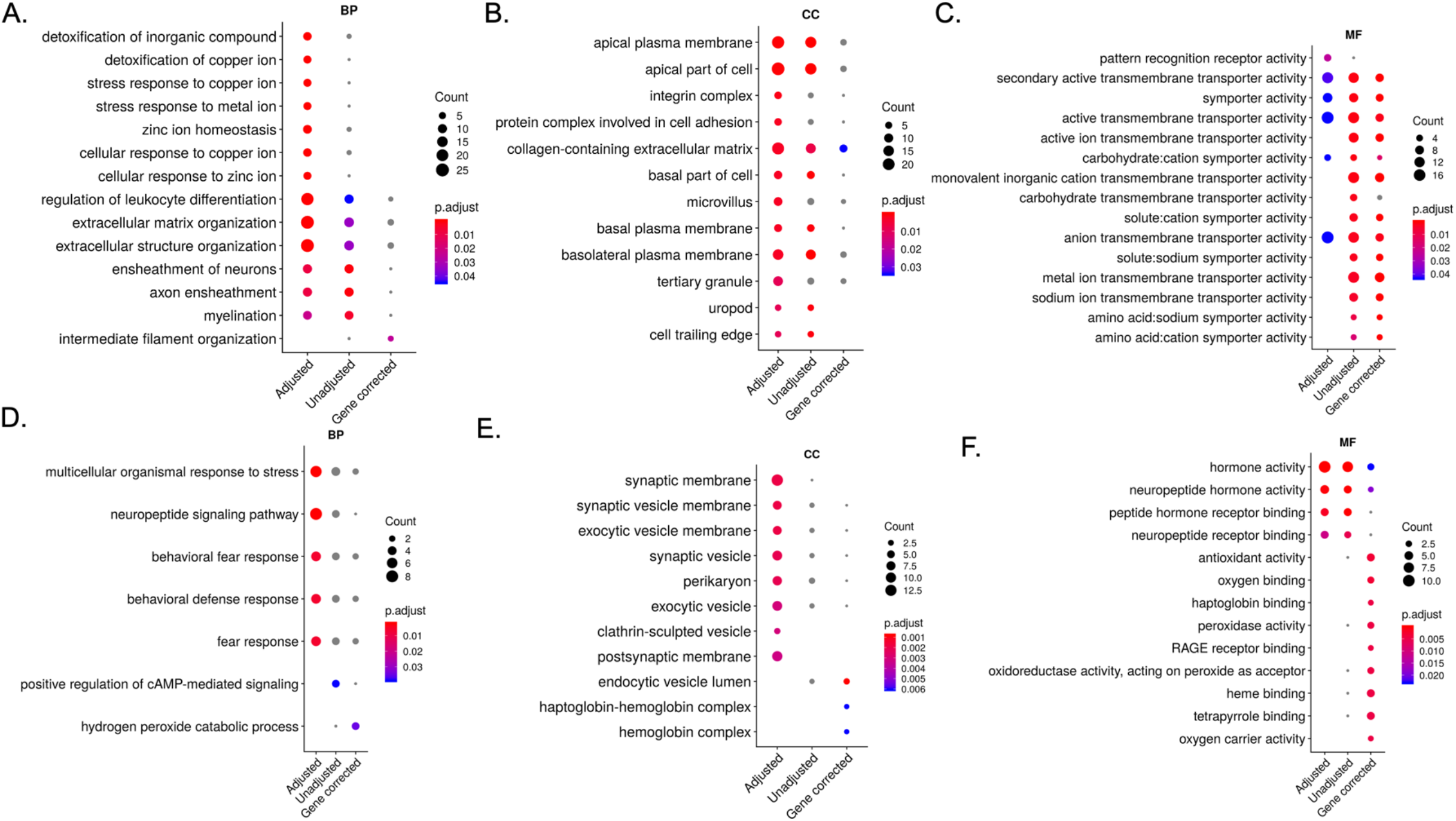
Comparison of GO enriched biological pathways before and after adjusting for the latent factor and after gene correction using a regression model.

**Fig. S6.**
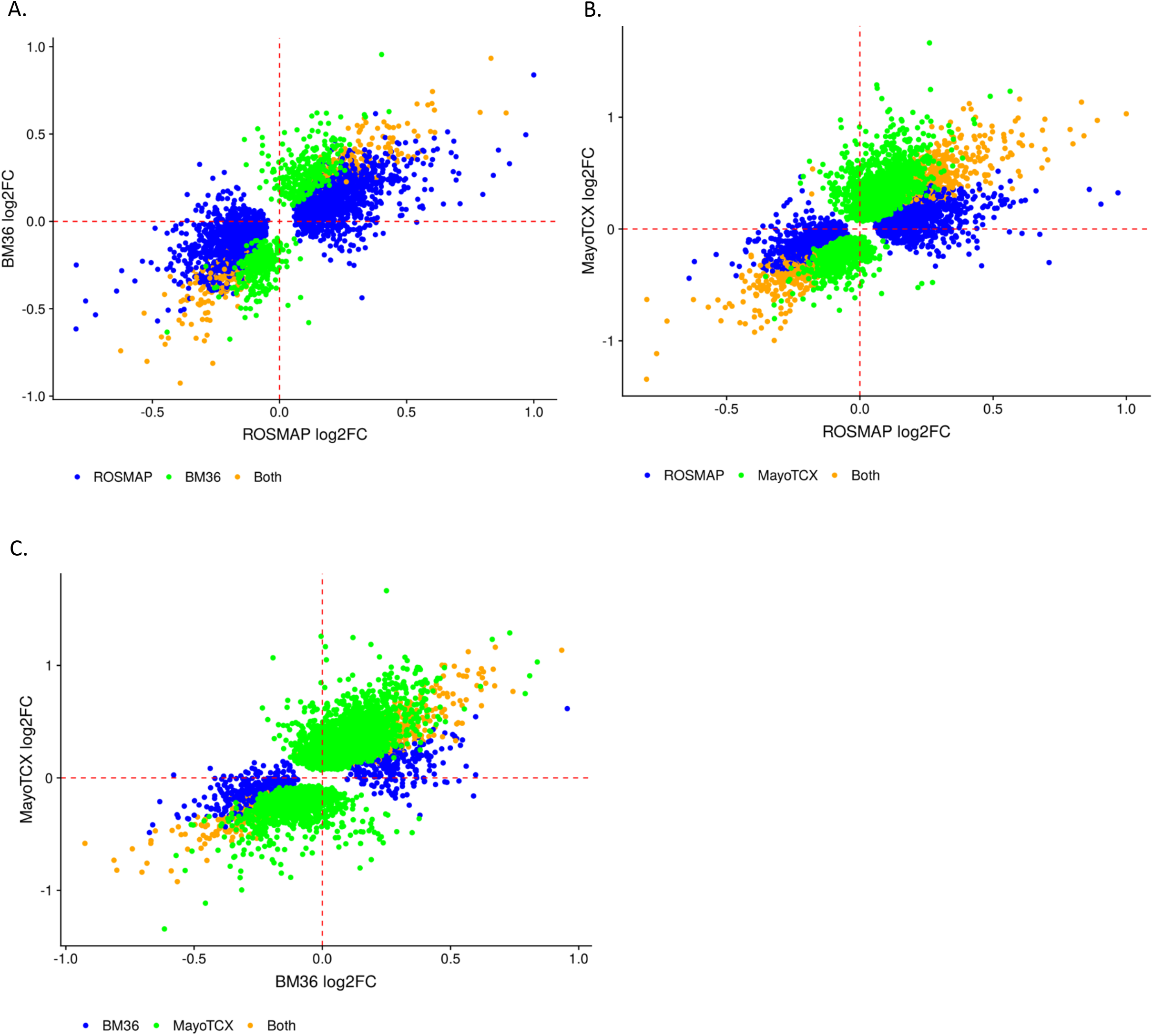
log2 fold change of DEGs comaprison after adjusting for the latent factor. DEGs have the same direction of change across studies after adjusting for the latent factor. DEGs which are signifcant in both studies are colored in orange.

